# Integration of pH control into Chi.Bio reactors and demonstration with small-scale enzymatic poly(ethylene terephthalate) hydrolysis

**DOI:** 10.1101/2024.03.03.582641

**Authors:** Mackenzie C.R. Denton, Natasha P. Murphy, Brenna Norton-Baker, Mauro Lua, Harrison Steel, Gregg T. Beckham

## Abstract

Small-scale bioreactors that are affordable and accessible would be of major benefit to the research community. In previous work, an open-source, automated bioreactor system was designed to operate up to the 30 mL scale with online optical monitoring, stirring, and temperature control, and this system, dubbed Chi.Bio, is now commercially available at a cost that is typically 1-2 orders of magnitude less than commercial bioreactors. In this work, we further expand the capabilities of the Chi.Bio system by enabling continuous pH monitoring and control through hardware and software modifications. For hardware modifications, we sourced low-cost, commercial pH circuits and made straightforward modifications to the Chi.Bio head plate to enable continuous pH monitoring. For software integration, we introduced closed-loop feedback control of the pH measured inside the Chi.Bio reactors and integrated a pH-control module into the existing Chi.Bio user interface. We demonstrated the utility of pH control through the small-scale depolymerization of the synthetic polyester, poly(ethylene terephthalate) (PET), using a benchmark cutinase enzyme, and compared this to 250 mL bioreactor hydrolysis reactions. The results in terms of PET conversion and rate, measured both by base addition and product release profiles, are statistically equivalent, with the Chi.Bio system allowing for a 20-fold reduction of purified enzyme required relative to the 250 mL bioreactor setup. Through inexpensive modifications, the ability to conduct pH control in Chi.Bio reactors widens the potential slate of biochemical reactions and biological cultivations for study in this system, and may also be adapted for use in other bioreactor platforms.

## Introduction

There is a major global effort underway to automate, down-scale, and democratize biological research. As a key component of these efforts, both commercial and open-source equipment are being developed for microbial cultivation and execution of biochemical reactions in a miniaturized context with online monitoring, with the intent to greatly accelerate the rate of data generation at lower cost, in reduced physical space, and with lower materials consumption (1-6). Moreover, to make biological research more accessible, there is a substantial drive for open-source hardware and software in biological research that often follows a do-it-yourself (DIY) model, enabled by the assembly of low-cost, off-the-shelf components (7-9). Of particular interest for biotechnological applications, commercial bioreactors with built-in control systems are usually quite costly, restricting their purchase primarily to companies and well-funded research laboratories. As a result, there have been many efforts to enable greater access to bioreactor hardware and software that, taken together, can result in the same or higher throughput than commercial bioreactors at 1-2 orders of magnitude cheaper, and using open-source software and DIY components (10-17).

Of relevance to the current work, the Chi.Bio system was introduced by Steel *et al*. in 2020 as an open-source bioreactor system that enables continuous bioprocess monitoring, spectrometry, light outputs, among other features (16). Steel *et al*. demonstrated the use of the parallelized Chi.Bio system in assays for cell growth, the formation of biofilms, control over optogenetics systems, and the simultaneous readout of orthogonal fluorescent protein signals. Conveniently, the Chi.Bio system is available both for purchase as parts that can be assembled, or the entire unit is available for purchase, which at the time of writing is for $990 for a control computer, reactor, and pump board. For systems like Chi.Bio, which can conduct both biochemical reactions and whole-cell cultivations, the ability to both continuously monitor and control the pH could expand the utility of this system to experimental systems where substrates or products that modify the pH of an aqueous medium dynamically vary.

To that end, here we integrate pH control into the Chi.Bio system through the modification of both the hardware, using off-the-shelf components, and through modifications to the open-source Chi.Bio software. We validate the ability of the Chi.Bio system to monitor and control pH using enzymatic hydrolysis of poly(ethylene terephthalate) (PET) as a demonstration application, and compare this to enzyme performance in a commercial 1 L bioreactor with an initial working volume of 250 mL. Overall, this work has a wide variety of potential applications across any process development involving pH-controlled biocatalytic reactions.

## Results

### Hardware integration of pH control functionality to Chi.Bio reactors

To realize a platform that enables pH monitoring and continuous pH control by acid or base addition, we modified the existing Chi.Bio system hardware (Labmaker). A list of all individual hardware components of the pH control module can be found in the Supplementary Information (**SI, Table 1**). The modifications comprise a customized head plate 3D-printed using a Stratasys Fortus 450MC printer with a 6.32 mm diameter port for insertion of a pH probe and a 1.5 mm diameter cut-out for an injector needle (**CAD file, SI**). A single silicon tubing line was connected to the injector needle, which inserts through the needle port of the customized head plate into the Chi.Bio reactor to provide a physical link for chemical addition back to the peristaltic pump board (**Figure 1A**). An off-the-shelf pH probe (ThermoScientific) was inserted centrally into the reactor via the headplate to enable real-time pH monitoring (**Figure 1B**). The pH probe physically connects to the pH circuit, built on a low-cost, easy-to-assemble breadboard integrating six pH Atlas-Scientific circuits to allow for the connection of up to six individual pH probes for the execution of up to six Chi.Bio reactors in parallel (**Figure 1C**).

**Table 1.** Yield quantification measured by mass loss and UPLC for PET degradation in Chi.Bio and Applikon bioreactors.

|  | Chi.Bio reactors | Applikon reactors |
| --- | --- | --- |
| Initial PET mass (g) | 1.2 | 25 |
| Final PET mass (g) | 0.089 ± 0.16 | 1.1 ± 0.50 |
| Extent of conversion by mass (%) | 92.6 ± 1.67 | 95.5 ± 2.02 |
| Extent of conversion by UPLC (%) | 94.3 ± 12.8 | 92.0 ± 16.4 |

**Figure 1.**
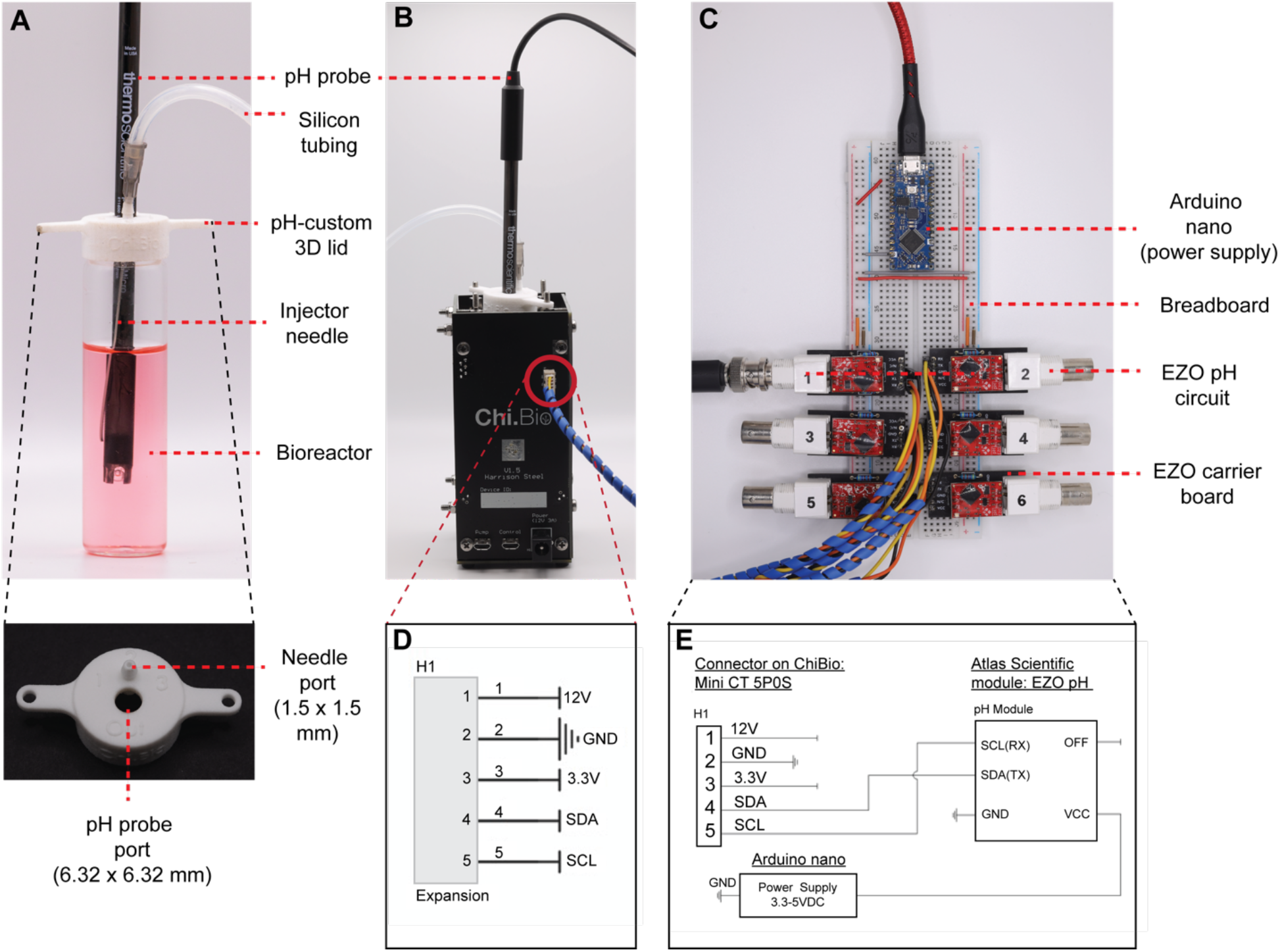
The hardware system for pH control integrated into Chi.Bio. The hardware modifications comprise an (**A**-**B**) off-the-shelf pH probe sourced from ThermoScientific (Orion™ Economy Series pH Combination Electrode #911600), inserted through a port in the custom 3D-printed head plate, the existing Chi.Bio peristaltic pump board (not shown), 2.5 mm x 1.0 mm silicon laboratory tubing (AlteSil High Strength Tubing, Altec, #01-93-1416), connected to an Air-Tite™ Premium Hypodermic Lab/Vet Use needle, and (**C**) an off-the-shelf Atlas Scientific pH circuit (EZO™ pH Circuit #EZO-pH) integrated to an Electrically Isolated EZO™ Carrier Board (#ISCCB-2). (**D**) Wiring diagram of expansion of the Chi.Bio Main Unit with the expansion header highlighted by red circle. (**E**) Wiring diagram of the Atlas Scientific EZO pH Module and ChiBio interface where all ground pins are connected to common ground. The 3.3V, 12V, and OFF pins are not connected.

For the circuitry integration, the original Chi.Bio reactor system includes an expansion header on the Main Reactor unit as shown circled in **Figure 1B**. The expansion header provides positive voltage, ground, and data connections to interface with the Chi.Bio controller (**Figure 1D**). Since data transmission occurs over the SDA (4) and SCL (5) pins using the I2C communication protocol, any I2C device can be implemented over this interface. Connection to this header was made using a 5-position mini-CT connector (Digikey). The EZO pH Circuit and Electrically Isolated EZO Carrier Board (Atlas Scientific) were selected as the pH transmitter using an off-the-shelf pH probe sourced from (ThermoScientific). The Carrier Board was connected to the Chi.Bio main reactor according to the wiring diagram in **Figure 1E**, where RX and TX are equivalently SCL and SDA, respectively per the manufacturer documentation. Connections were made using a breadboard (Sparkfun) for rapid prototyping. The 12V pin could be used to power the pH circuit if a voltage regulator was included. Instead, for simplicity a voltage regulator was not implemented, and a separate power supply (Arduino Nano) was used to provide 3.3V to the EZO Carrier Board to ensure the combined setup did not draw excessive current from the Chi.Bio 3.3V power rail. To turn off the EZO Carrier Board, the OFF pin can optionally be connected to GND. In this implementation, the OFF pin was left unconnected. Before installing the EZO pH Circuit in the system, the EZO was switched from serial communication to the I2C protocol by following the protocol selection guidance in the Atlas Scientific EZO documentation. The source files for all hardware pieces are available in the Beckham-lab GitHub (*vide infra*).

### Software integration of pH control functionality to Chi.Bio reactors

To achieve stable pH control in the Chi.Bio system, the pH of the reactor is compared to a target pH value, up to a set tolerance, and this is repeated at regular time intervals that we refer to as the cycle time. If the pH is outside of the set range, a fixed volume of acid or base is added to the system every 90 seconds to bring the pH back within the set limits. The cycle time is a user-defined quantity to allow for experiments to incorporate a variety of neutralizing agents and reaction conditions. In the following paragraphs, we discuss the details of how this feedback mechanism is incorporated into the existing Chi.Bio software.

The existing Chi.Bio system provides a user interface built in HTML/JavaScript that is accessible from a web browser and enables the user to have real-time control and monitor the experiment from a connected PC or network (**Figure 2A**). Behind the user interface, the Atlas Scientific EZO pH circuit was integrated into the existing I^2^C digital bus of Chi.Bio (**Figure 2B**). The Chi.Bio user interface was edited in HTML/JavaScript to incorporate a pH-control module (**Figure 2C**). The real-time measured pH is output to the UI webserver every 30 seconds to display the live pH value to the user (**Figure 2C**). Several tunable parameters were incorporated into the pH-control module in the form of clickable buttons, namely three buttons to calibrate the pH probe to pH 4, 7, and 10, one to select the desired pump line connected to the neutralizing agent (options 1-4), one to specify acid or base as the input, and one to activate the specified pump for the purpose of testing the flow rate. Additionally, three custom-input boxes were added to allow for the specification of the target pH, tolerance, and closed-loop control cycle time (**Figure 2C**). Finally, to prevent reactor overflow, the user can specify the maximum volume of neutralizing agent to be added the reactor in mL, as a built-in safety feature.

**Figure 2.**
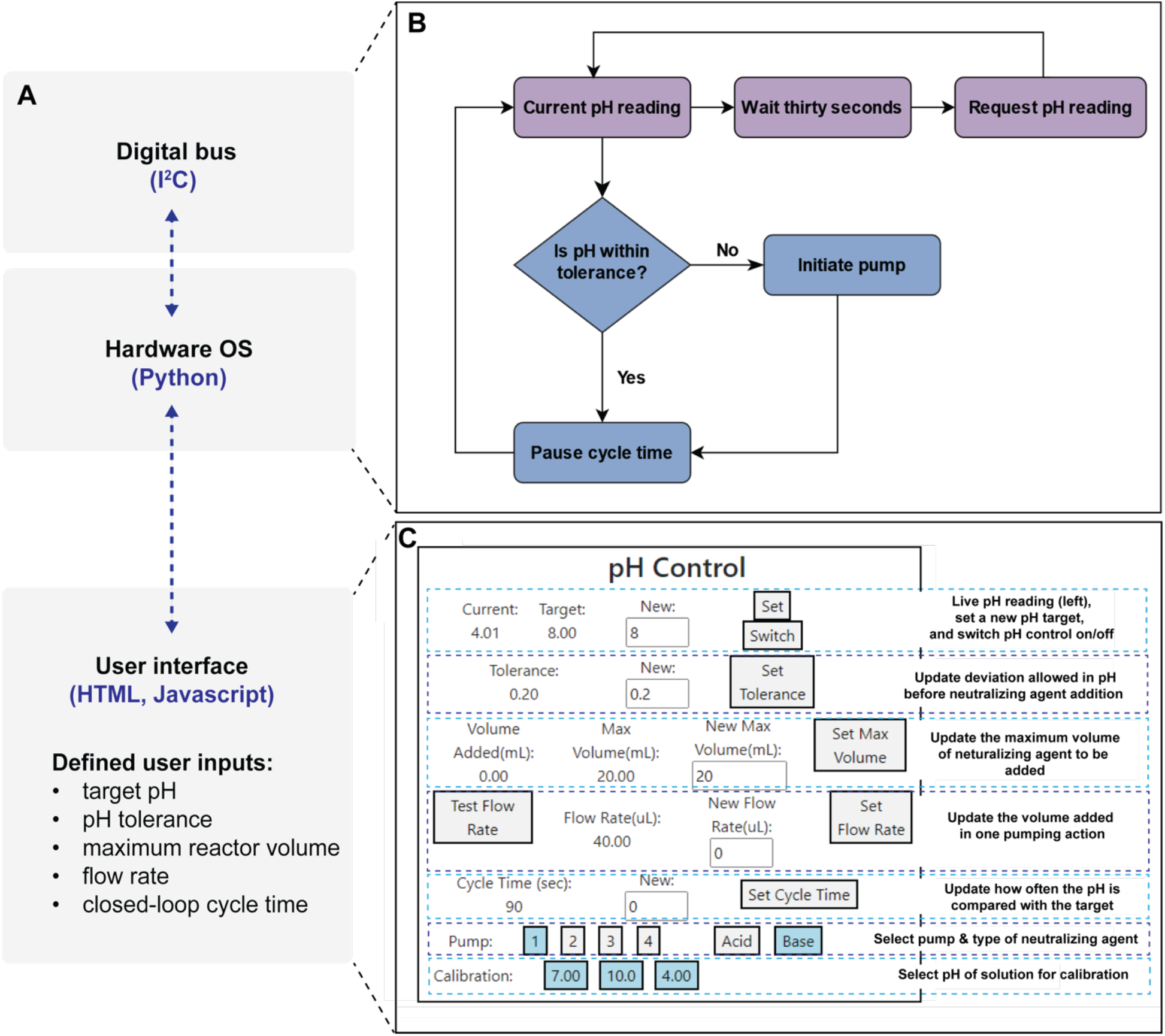
The software integration for pH control into Chi.Bio. (**A**) Overview of the Chi.Bio code that links the digital bus to the hardware OS and the live user interface. (**B**) Decision tree illustrating the function integrated into the digital bus and hardware OS to measure pH (upper horizontal flow cycle) and the function for the control of pH according to a predefined tolerance of a pH setpoint (lower cycle). (**C**) pH module embedded into the existing user Chi.Bio interface by modification to the HTML code.

The Python code linking the I^2^C digital bus to the real-time user interface was altered to introduce a closed-loop control on the pH inside the reactor as measured by the pH probe (**Figure 2B**). Specifically, a function was written to measure the pH every 30 seconds (**Figure 2B)**. A separate function was added to compare the current pH reading to a user-defined pH range (determined by pH ± tolerance) after the cycle time has elapsed. If the current pH reading lies outside the tolerated range, the pump is engaged to add a fixed volume of acid or base necessary to bring the pH within range (**Figure 2B**). The fixed volume is the volume added by a single on/off activation of the selected pump and input line. To determine the fixed volume, a ‘test flow rate’ button was included (**Figure 2C**). The volume added by the execution of this single on/off pump command corresponds to the fixed volume that will be added after each cycle time has elapsed. We manually determined the fixed volume to range between 30-45 ± 10 μL across individual Chi.Bio pumps for the set-up and application we detail below. The fixed volume achieved by a single on/off pump activation can additionally vary according to the specific neutralizing agent. For any given pump, input line, and neutralizing agent, it is recommended to repeat the measurement for an accurate distribution of fixed volume fluctuation, and to specify the fixed volume using the ‘set flow rate’ input box prior to each experiment. As our controller adds a fixed amount of liquid, in cases of large deviation from the pH set-point it might take multiple cycles of acid or base addition to return to the region of the set-point. This control approach is equivalent to an on-off controller (also known as *bang-bang*) with dead-zone, an effective and robust architecture often used in process control (18). Furthermore, this approach can be simplified and adapted for one-sided control (i.e. adding only acid or base during an experiment) when applied to regulate processes that drift toward high or low pH over time, as in the application detailed below.

The live pH readings are taken using the existing embed functionality in the Atlas Scientific pH circuit to correct for the temperature effect on pH. To ensure the pump adds the smallest possible volume of neutralizing agent, the pulse-width modulation was changed from 100% to 90%. Additionally, a custom function was written to block all other traffic on the I^2^C bus when the pumps are activated to ensure there is no delay in the off command being sent to the activated pump. The number of times the pump is activated in response to a change in pH is tracked and used to calculate the total volume of neutralizing agent added over the course of the experiment. For this calculation a “flow rate” needs to be measured and inputted by the user. Each experiment generates a comprehensive log file of the measured pH values and corresponding cumulative volume of neutralizing agent added.

### Applying the pH control functionality to enzymatic PET deconstruction

To demonstrate the utility of integrating pH control into the Chi.Bio platform, enzymatic hydrolysis reactions of poly(ethylene terephthalate) (PET) were carried out using the modified bioreactors relative to control reactions in 1-L scale Applikon bioreactors with a 250 mL working volume for the reaction, with the aim of demonstrating reactor-to-reactor reproducibility. Such control for the *in vitro* pH regulation of enzyme-mediated PET depolymerization reactions could enable rapid benchmarking and comparison of PET hydrolases for deconstruction across a wide range of industrially relevant reaction conditions and facilitate enzyme engineering (19-21). Namely, a major limitation of microplate format directed evolution and enzyme engineering campaigns is accumulation of the terephthalic acid product, which limits the extents of PET conversion at industrially relevant solid loadings (19). For example, phenol red dye-based indicator assays have been successfully applied for pH-sensitive monitoring of the soluble terephthalate and acidic oligomer products released by PETase activity, yet these approaches are also hampered by a lack of pH control (22).

Here, sodium hydroxide was used to neutralize the terephthalic acid product in real time of deconstruction of amorphous Goodfellow PET films by the ICCG variant of leaf-branch compost cutinase (hereafter LCC^ICCG^) (23), using the modified pH control functionality in the Chi.Bio system. Besides working volumes, the reaction conditions were identical between the Chi.Bio and Applikon reactor set-ups, where reactions were run in duplicate at 65°C with a 10% mass loading of Goodfellow PET film (ES301445), and an enzyme loading of 3 mg enzyme/g PET, in 100 mM sodium phosphate buffer at pH 8. The reactions achieved maintenance of pH control by applying a 0.1 pH unit tolerance range of the pH 8 target in the Chi.Bio reactors, and a 0.05 tolerance in the Applikon set-up (**Figure 3**). We chose 0.1 for the Chi.Bio pH regulation as this is the maximum limit of precision of the integrated ThermoScientific Orion™ Economy Series pH Combination Electrode. Across the two Chi.Bio reactors, the pH was maintained within 0.1 units of the target pH 8 for 78.7% and 80.2% of the total 72-hour reactions respectively, compared to 95.8 and 98.6% of the total reaction time in the Applikon reactors (**Figure 3, Datasets S1-S2**).

**Figure 3.**
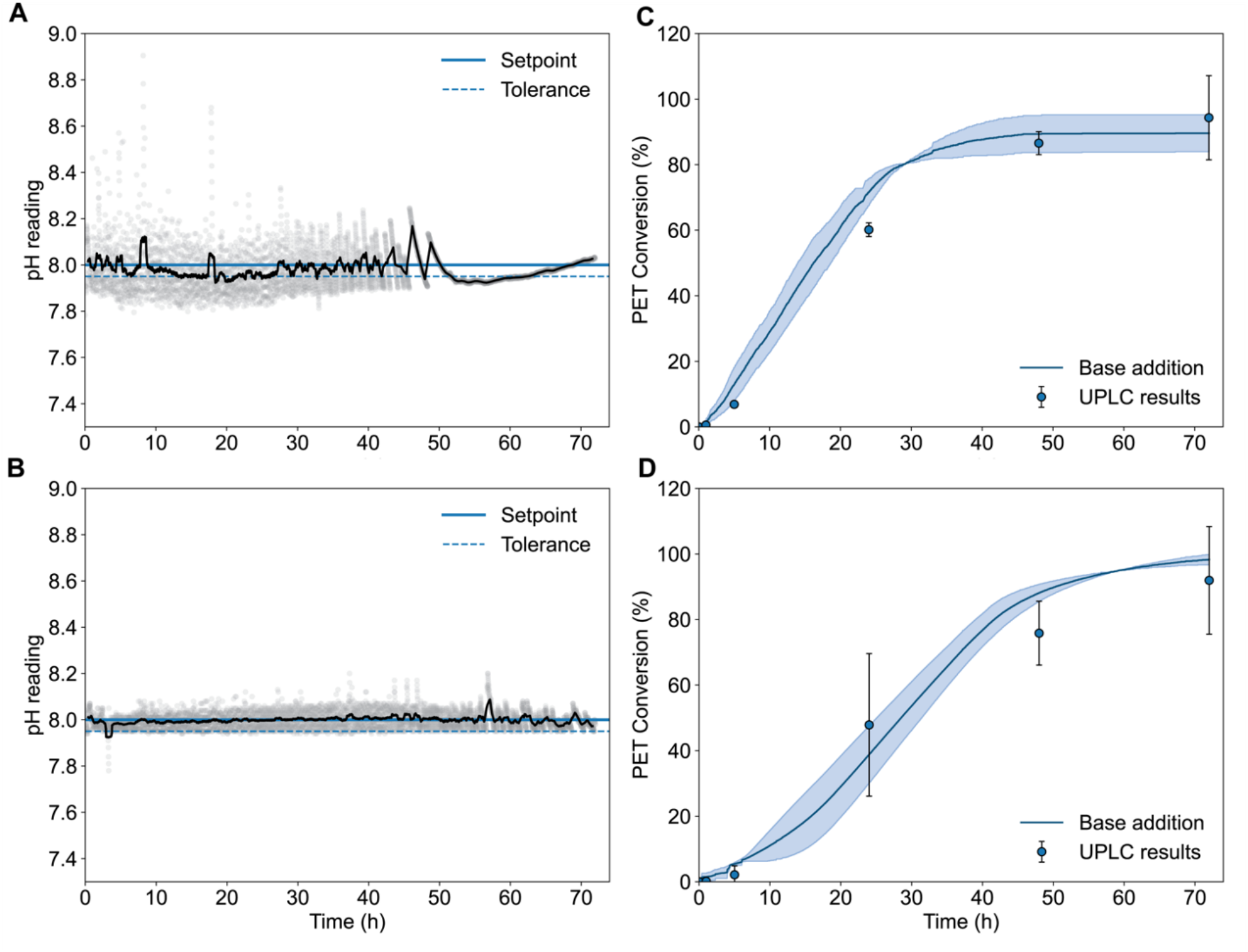
(**A**) Representative pH profiles with a rolling average pH calculated per 50 datapoints overlayed (black line) over all datapoints (light grey circles) in the Chi.Bio reactors, and (**B**) the Applikon bioreactors. The pH setpoint of 8 is shown as a solid blue line, and the respective tolerances (1 and 0.05 units below pH 8) as dashed-blue lines. (**C**) Extent of PET conversion as a function of time by PET hydrolase LCC^ICCG^ in the Chi.Bio reactors and (**D**) the Applikon bioreactors. The base addition extents of conversion (blue line) were calculated by sodium hydroxide addition, assuming that 2 mol of NaOH titrates 1 mol of TPA. The blue shaded zone represents the standard deviation of the calculated extent of PET conversion. Blue dots with error bars represent conversion based on UPLC quantification of TPA equivalents. The error bars denote the range of duplicate measurements.

After 72 hours, the contents of the reactors were filtered and dried to collect any unhydrolyzed PET. The final dried filtered solids were weighed to determine the final mass loss and used to calculate the extent of conversion of 92.6 ± 1.67% of PET in the Chi.Bio system, compared to 95.5 ± 2% in the Applikon reactors (**Table 1**). A conversion of 94.3 ± 12.8% was achieved as measured by ultra-high-performance liquid chromatography (UPLC) timepoint sampling in Chi.Bio by 72 hours, compared to a final 92 ± 16.4% in the Applikon bioreactors by 72 h respectively (**Figure 3**). At the 24 hr timepoint, the soluble monoacid product mono(2-hydroxyethyl) terephthalate (MHET) comprised 9.8% of the total aromatic product sum measured in the Chi.Bio reactors, which was all converted to TPA by 48 hr (**Dataset S3**). Comparatively, MHET was not detected in the Applikon reactor samples under these enzyme and solid loadings (**Dataset S4**), although MHET accumulation profiles can vary significantly according to reaction conditions and PET substrate selection (24).

While the final PET mass loss was similar in both systems (**Table 1**), a difference in the plateau of the rate of reaction was evident by comparison of the pH and base conversion profiles (**Figure 3**). These results suggest that pH tolerance values should be reported as part of standardization for comparison between studies of pH-controlled enzymatic PET deconstructions, as varying yield profiles may result from the application of different tolerances. Factors to explain the rate differences may include the differences in pH fluctuations, different surface area-to-volume ratios of the two reactor setups, or the different agitation schemes. In the Applikon reactors, constant agitation was applied at 400 rpm, compared to the existing stirring cycle time implemented in the Chi.Bio reactors which pauses every 60 seconds for data collection. Agitation rates of the enzyme and substrate, and corresponding shear stresses can also dictate enzyme stability over time in a reactor (25).

## Discussion

In this work, we implemented pH control software and hardware modifications into the Chi.Bio platform and demonstrated the utility of this modification for small-scale PET enzymatic hydrolysis reactions. We present this as a cost-effective platform that could be used to accelerate the evaluation of PET hydrolases in industrially relevant conditions. There has been a rapid expansion in the number of PET hydrolases and improved variants reported in the literature in the last 5 years alone (26-28). However, there is a need in the field to establish benchmarking of activity of the potentially process relevant PETases at industrially relevant loadings (19, 29, 30). In our results, we compared the robustness of the pH control in Chi.Bio to that implemented in Applikon bioreactors and established maintained conversion rates at the smaller scale, highlighting reproducibility in scaling-down base-controlled PETase reactions. This cost-effective system adds to a growing number of automated micro-reactor systems (31). It also opens opportunities to dissect a wide variety of polymer substrates in PETase deconstruction reactions without the requirement for gram to kilogram-scale substrate quantities and enables significant reductions in the purified enzyme required.

There are a variety of potential modifications that could further enhance the pH functionality we have integrated into the Chi.Bio system. For example, the pH probe that was integrated was the ThermoScientific Orion™ Economy Series pH Combination Electrode. Future modifications could incorporate a glass membrane pH probe, with an external glass electrode body, which may be better suited to repeated applications requiring high temperature or chemical resistance, and with greater precision than the 0.1 pH unit of precision of the existing probe. In the event of the integration of an alternative probe, the dimensions of the head plate could be correspondingly modified. The system utilizes the native Chi.Bio pump board, but could be swapped out for alternatives, such as a syringe pump, according to the sensitivity of regulation required. For microbial growth applications, the modified system is currently best suited to batch fermentations, or those requiring only a single-input line for neutralizing agent addition. Oxygen agitation for fermentation can be achieved with the existing Chi.Bio stirring function, and future modifications could also benefit from the integration of a dissolved oxygen stat module for aerobic cultivations. For the enzymatic depolymerization application shown here, the mass of the neutralizing agent and reactors before and after each enzymatic deconstruction was manually measured and tracked. The future integration of programmable micro-scales would allow for real-time monitoring of the neutralizing agent added and accurate quantification of reaction mass. Finally, the system could be applied in conjunction with robotic systems. Similar studies have integrated the original Chi.Bio platform to the low-cost Opentrons robotic liquid-handling systems for alternative applications (32).

### Conclusions

Overall, we have demonstrated the successful enzymatic deconstruction of PET with pH control in scaled-down PETase reactions. The PETase reactions demonstrated are a single application of the pH module we have integrated into the Chi.Bio platform. The functionality implemented should be broadly useful for a wide variety of biotechnological, biochemical, and synthetic biology applications including multi-enzymatic cascade reactions and pH-stat controlled growth of engineered strains of micro-organisms. The full range of biologically accessible pH values with corresponding control can be utilized for these applications using the Chi.Bio platform.

### Experimental procedures

#### Protein expression and purification

LCC^ICCG^ DNA was synthesized (Twist Bioscience) and cloned into a pET-21b(+) expression vector (EMD Biosciences) as previously described (24). The plasmid was transformed into OverExpress *Escherichia coli* C41 (DE3) (Lucigen) cells, plated on lysogeny broth (LB)-agar plates containing 100 μg/mL ampicillin (Amp), and incubated at 37 °C overnight. A single colony from transformation was inoculated into a starter culture of LB liquid media containing 100 μg/mL Amp and cultures were grown at 37 °C, 250 rpm overnight. The starter culture was inoculated at a 100-fold dilution in 2× YT media containing 100 μg/mL Amp and grown at 37 °C to OD_600_ = 0.6–0.8. Protein expression was induced by the addition of isopropyl β-D-1-thiogalactopyranoside (IPTG) at 1 mM. Cells were maintained at 18 °C, 225 rpm for 20 h following induction, harvested by centrifugation, and stored at −80 °C until purification. Harvested cells were resuspended in lysis buffer (20 mM Tris pH 8, 10 mM imidazole, 300 mM NaCl, 1 mg/mL lysozyme, 50 μg/mL DNAase) and subjected to sonication (QSonica Q700). The resulting lysate was clarified by centrifugation at 40,000 x *g* for 40 min at 4 °C. The clarified lysate was applied to a 25 mL HisTrap HP (Cytiva) column linked to a ÄKTA Pure chromatography system (Cytiva) and eluted with elution buffer (20 mM Tris pH 8, 500 mM imidazole, 300 mM NaCl) over a 2 CV gradient. The fractions containing the protein were dialyzed overnight into a 20 mM Tris, pH 8, 300 mM NaCl buffer and the protein purity confirmed by SDS-PAGE. The concentration was determined by 280 nm absorbance readings and calculated using an extinction coefficient of 37150 M^-1^ cm^-1^.

### Chi.Bio bioreactor PET deconstruction

Bioreactor hydrolysis reactions were performed in the Chi.Bio reactors with a reaction volume of 12 mL, with Goodfellow PET film (ES301445) cut into 1 cm x 1 cm squares as the substrate. For the reactions, 1.2 g of PET substrate (1.38 mL volume based on density of PET) was added to 100 mM phosphate pH 8 assay buffer at a final volume of 12 mL and equilibrated to 65 °C with a stirring rate set to 0.2. The reactions were initiated by the addition of 0.77 mL of 4.7 mg/mL LCC-ICCG for a final enzyme loading of 3 mg/g PET. Reactions proceeded for 72 h and were maintained at pH 8 with 1 M NaOH addition using the integrated pH functionality. Sample volumes of 0.1 mL were removed at designated time points, quenched with an equal volume of methanol, stored, and filtered. At the end of the reaction time course, the remaining substrate was collected by filtration through Whatman grade 2 filter paper (Cytiva) and a Büchner funnel. The filters were preweighed, and the filters with PET were dried for 3 days at 40 °C under vacuum before the final mass of residual PET was calculated.

### Applikon bioreactor PET deconstruction

Bioreactor hydrolysis reactions were performed at 0.25 L scale in duplicate in 1 L glass bioreactors (Applikon Biotechnology), which included two Rushton impellers in the stirrer shaft below the 200 mL line. The substrate used was Goodfellow PET film (ES301445) cut into 2.5 cm x 2.5 cm squares. For the reactions, 25 g of PET substrate (18 mL volume based on density of PET) was added to 100 mM phosphate pH 8 assay buffer at a final volume of 0.25 L and equilibrated to 65 °C with stirring at 400 rpm. The reactions were initiated by the addition of 16 mL of 4.7 mg/mL LCC-ICCG for a final enzyme loading of 3 mg/g PET. Reactions proceeded for 72 h and were maintained at pH 8 with 4 M NaOH addition using a peristaltic pump controlled by an in-control module (Applikon Biotechnology). Sample volumes of 0.5 mL were removed at designated time points, quenched, stored, and filtered. At the end of the reaction time course, the remaining substrate was collected by filtration through Whatman grade 2 filter paper (Cytiva) and a Büchner funnel. The filters were preweighed, and the filters with PET were dried for 3 days at 40 °C under vacuum before the final mass of residual PET was calculated.

### UPLC quantification

Analysis of aromatic products MHET, BHET and TPA was performed by ultra-high performance liquid chromatography (UHPLC) as previously described (21). Briefly, samples were injected onto a Zorbax Eclipse Plus C18 Rapid Resolution HD column and separation was achieved using a mobile phase gradient of 20 mM phosphoric acid and methanol. Diode array detection (DAD) was utilized for quantitation for the analytes of interest using a wavelength of 240 nm.

## Data availability

Code for the pH control for Chi.Bio and the source files for all hardware pieces are available in the Beckham-lab GitHub at https://github.com/beckham-lab/Chi.Bio.pH.

## Supporting Information

This article contains supporting information.

## Acknowledgements

We thank the members of the BOTTLE Consortium and the Centre for Enzyme Innovation for helpful discussions. We thank Christine Singer and Davinia Salvachúa for help with the Applikon bioreactor setup. We thank Hannah Alt and Kelsey J. Ramirez for assistance with analytics.

## Author contributions

Conceptualization: MCRD, NPM, GTB; Resources: ML, HS; Software: MCRD; Funding acquisition: GTB; Investigation: MCRD, NPM, BNB; Visualization: MCRD, NPM; Methodology: MCRD, NPM; Writing – original draft: MCRD, NPM, GTB; Writing – review and editing: all authors.

## Funding and additional information

Funding for MCRD, NPM, and GTB was provided by the US Department of Energy (DOE), Office of Energy Efficiency and Renewable Energy, Advanced Materials and Manufacturing Technologies Office (AMMTO) and Bioenergy Technologies Office (BETO). This work was performed as part of the Bio-Optimized Technologies to keep Thermoplastics out of Landfills and the Environment (BOTTLE) Consortium and was supported by AMMTO and BETO under contract no. DE-AC36-08GO28308 with the National Renewable Energy Laboratory (NREL), operated by Alliance for Sustainable Energy, LLC. This material is also based upon work from BNB and GTB supported by the U.S. Department of Energy, Office of Science, Office of Biological and Environmental Research, Genomic Science Program under Award Number DE-SC0022024. HS was supported in part by the UK’s Engineering and Physical Sciences Research Council (EPSRC) project EP/W000326/1. The views expressed in the article do not necessarily represent the views of the DOE or the U.S. Government. The U.S. Government retains and the publisher, by accepting the article for publication, acknowledges that the U.S. Government retains a nonexclusive, paid-up, irrevocable, worldwide license to publish or reproduce the published form of this work, or allow others to do so, for U.S. Government purposes.

## Supplementary Information

**Supplementary Table 1 (S1):**
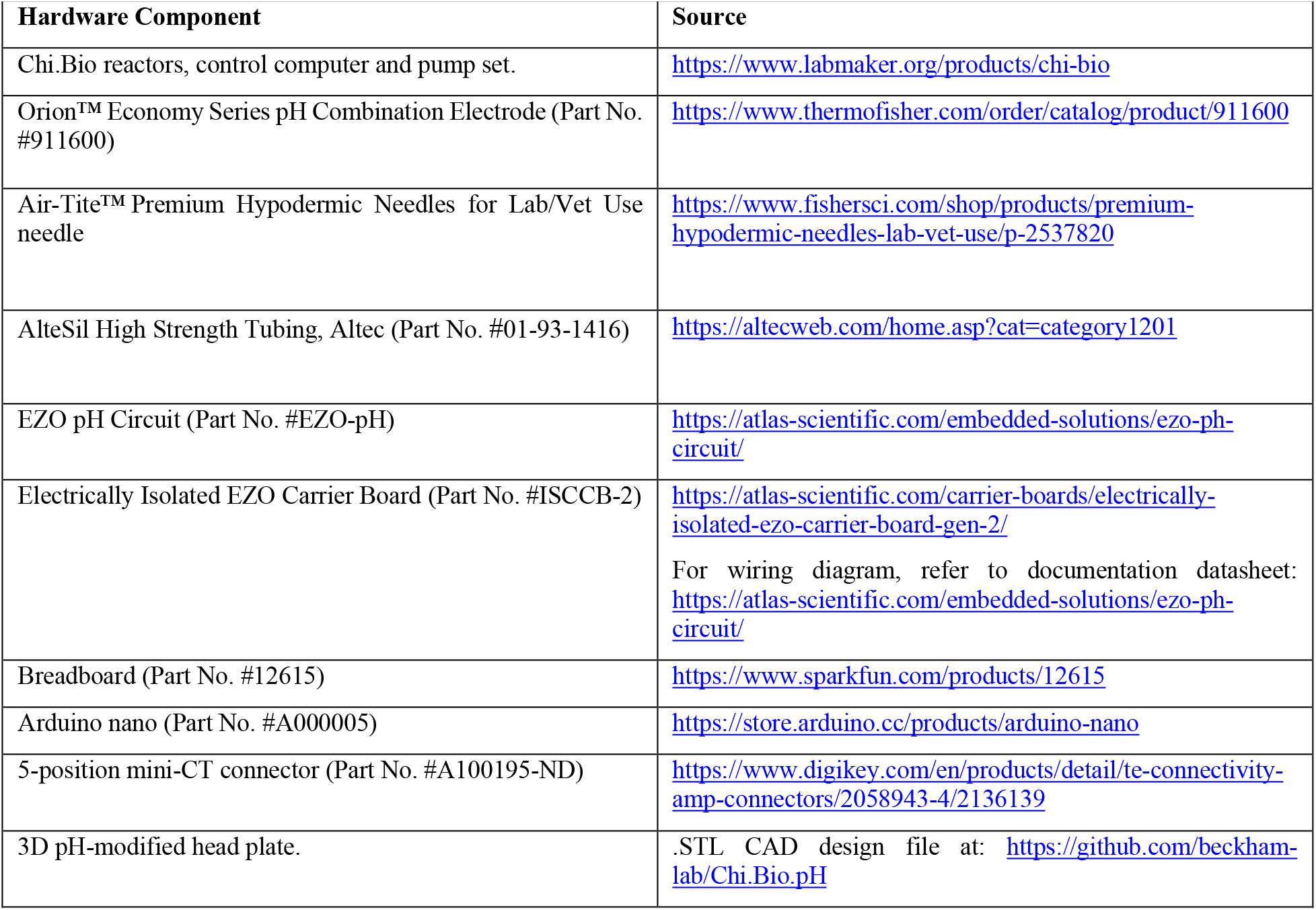
Hardware components for the pH system integrated into the Chi.Bio platform.

## Supplementary Files

- https://github.com/beckham-lab/Chi.Bio.pH (app.py, HTMLScripts.js, index.html, README.txt, pH_Probe_Cap.STL)
- Supplementary Datasets S1-S4 (Excel File) S1: Base profile and base conversion dataset for enzymatic PET deconstructions in Chi.Bio bioreactors. S2: Base profile and base conversion dataset for enzymatic PET deconstructions in Applikon bioreactors. S3: UPLC datasets for enzymatic PET deconstructions in Chi.Bio bioreactors. S4: UPLC datasets for enzymatic PET deconstructions in Applikon bioreactors.

## References

1. Betts JI, Baganz F. 2006. Miniature bioreactors: current practices and future opportunities. Microb. Cell Factories 5: 21.

2. King RD, Rowland J, Oliver SG, Young M, Aubrey W, et al. 2009. The automation of science. Science 324: 85–89.

3. Klein T, Schneider K, Heinzle E. 2013. A system of miniaturized stirred bioreactors for parallel continuous cultivation of yeast with online measurement of dissolved oxygen and off-gas. Biotechnol. Bioeng. 110: 535–42.

4. Tajsoleiman T, Mears L, Krühne U, Gernaey KV, Cornelissen S. 2019. An industrial perspective on scale-down challenges using miniaturized bioreactors. Trends Biotechnol. 37: 697–706.

5. Martin HG, Radivojevic T, Zucker J, Bouchard K, Sustarich J, et al. 2023. Perspectives for self-driving labs in synthetic biology. Curr. Opinion Biotechnol. 79: 102881.

6. Bajić M, Khiawjan S, Hilton ST, Lye GJ, Marques MPC, Szita N. 2024. A paradigm shift for biocatalytic microreactors: Decoupling application from reactor design. Biochem. Eng. J.: 109260.

7. Wenzel T. 2023. Open hardware: From DIY trend to global transformation in access to laboratory equipment. PLoS Biology 21: e3001931.

8. Castro-García JA, Molina-Cantero AJ, Merino-Monge M, Gómez-González IM. 2019. An open-source hardware acquisition platform for physiological measurements. IEEE Sensors Journal 19: 11526–34.

9. Pearce JM. 2012. Building research equipment with free, open-source hardware. Science 337: 1303–04.

10. Miller AW, Befort C, Kerr EO, Dunham MJ. 2013. Design and use of multiplexed chemostat arrays. JoVE: e50262.

11. Toprak E, Veres A, Yildiz S, Pedraza JM, Chait R, et al. 2013. Building a morbidostat: an automated continuous-culture device for studying bacterial drug resistance under dynamically sustained drug inhibition. Nature Protocols 8: 555–67.

12. Takahashi CN, Miller AW, Ekness F, Dunham MJ, Klavins E. 2015. A low cost, customizable turbidostat for use in synthetic circuit characterization. ACS Syn. Biol. 4: 32–38.

13. Wong BG, Mancuso CP, Kiriakov S, Bashor CJ, Khalil AS. 2018. Precise, automated control of conditions for high-throughput growth of yeast and bacteria with eVOLVER. Nature Biotechnol. 36: 614–23.

14. Bergenholm D, Liu G, Hansson D, Nielsen J. 2019. Construction of mini-chemostats for high-throughput strain characterization. Biotechnol. Bioeng. 116: 1029–38.

15. McGeachy AM, Meacham ZA, Ingolia NT. 2019. An accessible continuous-culture turbidostat for pooled analysis of complex libraries. ACS Syn. Biol. 8: 844–56.

16. Steel H, Habgood R, Kelly CL, Papachristodoulou A. 2020. In situ characterisation and manipulation of biological systems with Chi. Bio. PLoS Biology 18: e3000794.

17. Gopalakrishnan V, Crozier D, Card KJ, Chick LD, Krishnan NP, et al. 2022. A low-cost, open-source evolutionary bioreactor and its educational use. eLife 11: e83067.

18. Janert PK. 2013. Feedback Control for Computer Systems. Sebastopol: O’Reilly Media, Inc.

19. Arnal G, Anglade J, Gavalda S, Tournier V, Chabot N, et al. 2023. Assessment of four engineered PET degrading enzymes considering large-scale industrial applications. ACS Catal. 13: 13156–66.

20. Tournier V, Duquesne S, Guillamot F, Cramail H, Taton D, et al. 2023. Enzymes’ power for plastics degradation. Chem. Rev. 123: 5612–701.

21. Erickson E, Gado JE, Avilán L, Bratti F, Brizendine RK, et al. 2022. Sourcing thermotolerant poly (ethylene terephthalate) hydrolase scaffolds from natural diversity. Nature Comm. 13: 7850.

22. Beech JL, Clare R, Kincannon WM, Erickson E, McGeehan JE, et al. 2022. A flexible kinetic assay efficiently sorts prospective biocatalysts for PET plastic subunit hydrolysis. RSC Adv. 12: 8119–30.

23. Tournier V, Topham C, Gilles A, David B, Folgoas C, et al. 2020. An engineered PET depolymerase to break down and recycle plastic bottles. Nature 580: 216–19.

24. Brizendine RK, Erickson E, Haugen SJ, Ramirez KJ, Miscall J, et al. 2022. Particle size reduction of poly (ethylene terephthalate) increases the rate of enzymatic depolymerization but does not increase the overall conversion extent. ACS Sust. Chem. Eng. 10: 9131–40.

25. Hayes HC, Luk LYP. 2023. Investigating the effects of cyclic topology on the performance of a plastic degrading enzyme for polyethylene terephthalate degradation. Sci Rep 13: 1267.

26. Cui Y, Chen Y, Liu X, Dong S, Tian Ye, et al. 2021. Computational redesign of a PETase for plastic biodegradation under ambient condition by the GRAPE strategy. ACS Catal. 11: 1340–50.

27. Erickson E, Shakespeare TJ, Bratti F, Buss BL, Graham R, et al. 2021. Comparative performance of PETase as a function of reaction conditions, substrate properties, and product accumulation. ChemSusChem 15: e202101932.

28. Son HF, Cho IJ, Joo S, Seo H, Sagong H-Y, et al. 2019. Rational protein engineering of thermo-stable PETase from Ideonella sakaiensis for highly efficient PET degradation. ACS Catal. 9: 3519–26.

29. Singh A, Rorrer NA, Nicholson SR, Erickson E, DesVeaux JS, et al. 2021. Techno-economic, life-cycle, and socioeconomic impact analysis of enzymatic recycling of poly (ethylene terephthalate). Joule 5: 2479–503.

30. Uekert T, Desveaux JS, Singh A, Nicholson SR, Lamers P, et al. 2022. Life cycle assessment of enzymatic poly(ethylene terephthalate) recycling. Green Chem. 24: 6531–43.

31. Fritzsche S, Tischer F, Peukert W, Castiglione K. 2023. You get what you screen for: a benchmark analysis of leaf branch compost cutinase variants for polyethylene terephthalate (PET) degradation. Reaction Chem. Eng. 8: 2156–69.

32. Bertaux F, Sosa-Carrillo S, Gross V, Fraisse A, Aditya C, et al. 2022. Enhancing bioreactor arrays for automated measurements and reactive control with ReacSight. Nature Comm. 13: 3363.

